# DINMC: A Deep Learning Framework for Interpretable Normative Model Construction and Pathological Brain Alteration Detection

**DOI:** 10.64898/2026.05.29.728652

**Authors:** Zewu Ge, Shui Liu, Weibei Dou

## Abstract

**Background and Objective:** Normative modeling is a key tool for understanding brain alterations in neurodegenerative diseases, such as cerebellar-type multiple system atrophy. However, existing methods lack interpretability and fail to capture clinically meaningful pathological changes. This study presents DINMC, a Deep Interpretable Normative Model Construction framework, which combines autoencoder-based learning with statistical hypothesis testing to better capture and interpret disease-specific neuroanatomical changes.

**Methods:** The DINMC framework constructs normative models using neuroimaging data from multi-site large healthy cohorts. It utilizes a U-shaped convolutional autoencoder to train these models, which are then applied to reconstruct brain features from both patients and healthy controls within the same study cohort. Pathological confidence values are derived by fusing original and deviation feature spaces, offering a measure of disease-related pathology reflected in each dimension of the features. The framework was validated through statistical analysis and prognostic classification and regression tasks.

**Results:** The pathological confidence provides valuable insights into the neuroanatomical regions most affected by the disease, as well as the correlation between changes in these regions and clinical assessment scales. Our optimal model outperform traditional methods in prognostic prediction tasks, with an AUC of 0.972 for classification tasks and an *R*^2^ of 0.432 for regression tasks.

**Conclusion:** DINMC provides a novel and interpretable framework for neuroimaging analysis. By combining deep learning and statistical hypothesis testing, this framework offers a unique solution to improving both the interpretability and performance of normative models in neuroimaging. The approach is scalable to other neuroimaging datasets, offering a versatile tool for broader biomedical applications.

## 1. Introduction

Normative models are standardized population-based frameworks designed to quantify biological variability within large healthy cohorts [1]. Analogous to growth charts in pediatric medicine, lifespan brain charts have been proposed to characterize normative trajectories of brain development and aging [2]. Statistical approaches for modeling population distributions and individual deviations have long been applied in biomedical research [3], although these methods were not explicitly framed as “normative models.” Unlike binary classifications of individuals as “normal” or “abnormal,” normative models explicitly characterize the distribution of biological measures within a healthy population [4], thereby enabling individualized deviation assessment [5, 6]. With the recent release of large-scale public magnetic resonance imaging (MRI) datasets of healthy individuals [7, 8, 9, 10], normative modeling has been increasingly applied to the diagnosis and personalized treatment of neurological and psychiatric disorders [11, 12, 13, 14].

The construction of robust normative models critically depends on high-quality reference data [15]. Although large multi-site datasets alleviate data scarcity, their effectiveness is challenged by substantial inter-site heterogeneity arising from protocol differences, scanner hardware, and acquisition parameters [16, 17]. ComBat, an empirical Bayesian harmonization technique originally developed for gene expression analysis [18], has been widely adopted to mitigate site effects in multi-site neuroimaging studies [19, 20]. Alternatively, some normative modeling approaches explicitly include site effects as covariates during model construction. However, such data-dependent correction strategies may inadvertently remove biologically meaningful signals, particularly when clinical covariates are intrinsically correlated with site effects [21]. As a result, aggressive site correction can obscure disease-related variability rather than clarify it.

A variety of methodological approaches have been proposed for normative model construction, including Gaussian process regression [3, 14], generalized additive models for location, scale, and shape (GAMLSS) [2, 15], autoencoder-based deep learning models [11, 22, 23], and other regression-based frameworks [21, 24]. Several of these methods have been integrated into open-source software platforms that leverage large healthy population datasets [25, 26]. Following normative model construction, deviation metrics—such as Z-scores—are typically derived and used as substitutes for original features in downstream analyses [1, 14]. While deviation features have been shown to enhance group-level statistical sensitivity and improve classification or prediction performance [27], the complementary information contained in the original features is often underutilized.

Moreover, many existing normative modeling frameworks suffer from limited clinical interpretability. Neurobiological features are inherently multidimensional [28], with spatially distributed patterns that encode clinically relevant pathological information [29]. However, current approaches typically either collapse multidimensional features into a single deviation metric or treat each dimension independently without a principled mechanism for identifying pathologically meaningful dimensions. Although covariates can be statistically adjusted, the biological interpretation of such adjustments remains unclear. Traditional clinical neuroimaging studies often rely on carefully matched pathological control groups to address confounding factors, but limited sample sizes constrain analyses to group-level hypothesis testing. Importantly, normative modeling is fundamentally a bottom-up, data-driven approach and cannot replace theory-driven hypothesis testing [17], particularly given the risk of introducing spurious biological signals during covariate correction. Consequently, there is a critical need for interpretable normative modeling frameworks that explicitly integrate statistical inference with deviation modeling to support biologically grounded and clinically meaningful conclusions.

Here, we propose a Deep Interpretable Normative Model Construction (DINMC) framework that integrates statistical hypothesis testing with autoencoder-based normative modeling. The framework begins by training a U-shaped convolutional autoencoder (UCAE) on a large-scale multi-site healthy cohort. The trained normative model learns the feature distribution patterns of the healthy population while simultaneously reconstructing features of patients to resemble those of matched healthy controls[23]. Building on this model, we introduce the concept of pathological confidence, which quantifies the degree to which features reflect genuine disease-related pathology. We hypothesize that when original features are pathologically relevant, the normative model will struggle to accurately reconstruct them, leading to the creation of deviation features that highlight pathological differences. By statistically comparing significant differences between the original and deviation feature spaces, we compute dimensional pathological confidence values, enabling precise identification of neuroanatomical regions that contribute to clinical abnormalities.

## 2. Materials and Methods

Fig. 1 illustrates the main modules and key concepts of the DINMC framework applied to real clinical datasets. The details of each part of Fig. 1 are presented throughout Sections 2.1–2.4.

**Figure 1.**
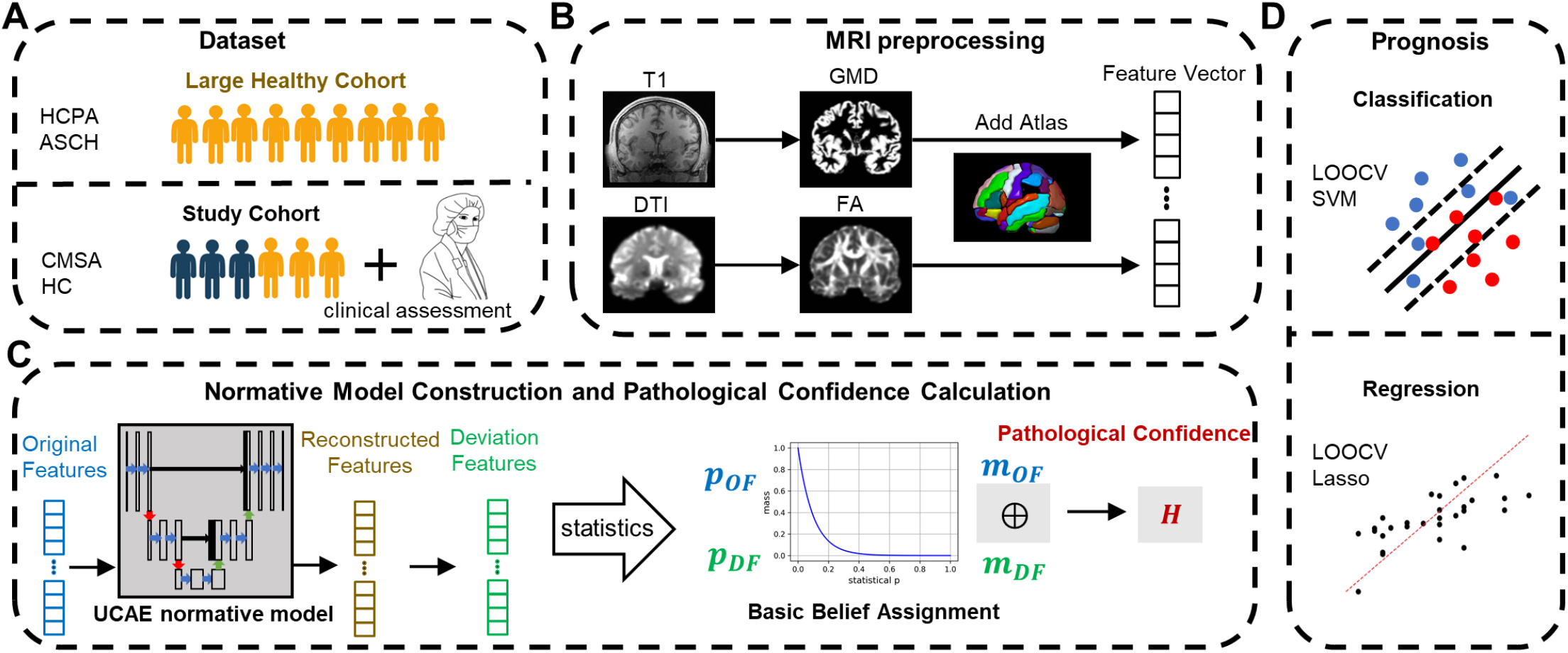
Overview of the proposed DINMC Framework. (A) Datasets used in this study included both large healthy cohorts and study cohorts (Human Connectome Project Aging, HCPA; AeroSpace Center Hospital, ASCH; cerebellar-type multiple system atrophy, CMSA; Healthy Controls, HC).(B) Multimodal MRI features were extracted through standardized processing: gray matter density (GMD) was derived from T1-weighted imaging while fractional anisotropy (FA) was calculated from diffusion tensor imaging (DTI). Regional feature averaging was performed using the brain atlas to obtain brain region-level features. (C) The framework builds a U-shaped convolutional autoencoder (UCAE) normative model using healthy control data to reconstruct original features and derive deviation features. These are then analyzed through statistical tests and fused to calculate pathological confidence (H) values using the Dempster-Shafer Evidence Theory. (D) Prognosis was conducted through leave-one-out cross-validation (LOOCV) using both support vector machine (SVM) classification and Lasso regression.

### 2.1. Datasets

This study utilized multimodal MRI data from three distinct sources. Two large-scale healthy cohorts were included: the Human Connectome Project Aging (HCPA)[10], the AeroSpace Center Hospital (ASCH) dataset. The third dataset was a clinical study cohort from Dongfang Hospital of Beijing University of Chinese Medicine, comprising 29 patients with cerebellar-type multiple system atrophy (CMSA) and 27 healthy controls (HC)[30]. Each patient was assessed by the Unified Multiple System Atrophy Rating Scale (UMSARS), which comprises Part I (UMSARS-I): a 12-item historical/activities-of-daily-living questionnaire and Part II (UMSARS-II): a 14-item clinician-rated motor examination; higher scores indicate greater disability. To ensure demographic comparability, healthy subjects from large-scale healthy cohorts were selectively matched to the clinical participants by age and gender. Table 1 shows the detailed demographics.

**Table 1:** Demographic and Clinical Characteristics of the Multimodal MRI Datasets.

| dataset | Age, years | Sex M/F | UMSARS-I | UMSARS-II |
| --- | --- | --- | --- | --- |
| HCPA | 56.29<sup>1</sup>±6.91<sup>2</sup> | 136/184 | NA<sup>3</sup> | NA |
| ASCH | 55.35<sup>1</sup>±5.59<sup>2</sup> | 94/101 | NA | NA |
| CMSA | 57.62<sup>1</sup>±6.00<sup>2</sup> | 18/11 | 17.34<sup>1</sup>±5.65<sup>2</sup> | 17.07<sup>1</sup>±5.66<sup>2</sup> |
| HC | 57.37<sup>1</sup>±5.64<sup>2</sup> | 11/16 | NA | NA |

UMSARS, Unified Multiple System Atrophy Rating Scale;
<sup>1</sup> represents average value;
<sup>2</sup> stands for standard deviation;
<sup>3</sup> represents Not Applicable.

### 2.2. MRI Acquisition and Preprocessing

The primary multimodal MRI modalities analyzed were T1-weighted imaging and diffusion tensor imaging (DTI). Detailed information about these datasets and their acquisition parameters is presented in the Table 2 and Table 3.

**Table 2:** T1-weighted imaging scan parameters for different datasets.

| Parameter | ASCH | HCPA | CMSA |
| --- | --- | --- | --- |
| Manufacturer | Siemens | Siemens | GE |
| Magnetic field strength (T) | 3 | 3 | 3 |
| Repetition time (ms) | 2530 | 2500 | 8172 |
| Echo time (ms) | 3.07 | 1.8/3.6/5.4/7.2 | 3.18 |
| Inversion time (ms) | 1100 | 1000 | 450 |
| Flip angle (°) | 12 | 8 | 12 |
| Voxel spacing (mm<sup>3</sup>) | 0.4316 × 0.4316 × 1 | 0.8 × 0.8 × 0.8 | 0.5 × 0.5 × 1 |
| Acquisition matrix | 512 × 512 × 19 | 320 × 300 × 208 | 512 × 512 × 18 |

**Table 3:** Diffusion tensor imaging (DTI) scan parameters for different datasets.

| Parameter | ASCH | HCPA | CMSA |
| --- | --- | --- | --- |
| Manufacturer | Siemens | Siemens | GE |
| Magnetic field strength (T) | 3 | 3 | 3 |
| Repetition time (ms) | 2700 | 3230 | 6000 |
| Echo time (ms) | 82 | 89.2 | 62.2 |
| Voxel spacing (mm<sup>3</sup>) | 2 × 2 × 2 | 1.5 × 1.5 × 1.5 | 1 × 1 × 3 |
| Acquisition matrix | 112 × 112 × 76 | 140 × 140 × 92 | 256 × 256 × 50 |
| Directions | 128 | 99 | 64 |
| b-values (s/mm<sup>2</sup>) | 1000 / 2000 | 1500 / 3000 | 1000 |

All T1-weighted and DTI data were processed through standardized pipelines. T1 preprocessing was performed in SPM12 software[31] in MATLAB 2023b (MathWorks, Natick, MA, United States) for tissue segmentation by a posteriori maximum estimation within a Gaussian mixture model framework, producing gray matter density (GMD) maps. DTI preprocessing was performed in FSL 6.0’s diffusion toolbox (FMRIB, Oxford)[32], implementing brain extraction, eddy current correction, and tensor fitting to generate fractional anisotropy (FA) maps. GMD and FA feature maps were spatially normalized to the MNI152 template at 1*mm*^3^ and 2*mm*^3^ isotropic resolution using FSL[32], respectively.

Multimodal feature extraction integrated three different brain parcellation atlas, including Automated Anatomical Labeling (AAL116) atlas[33],the Atlas of Intrinsic Connectivity of Homotopic Areas (AICHA384)[34] and the Shen268 atlas[35]. For each participant, region-specific features were calculated by averaging voxel values within each ROI defined by brain atlas, generating distinct multidimensional feature vectors corresponding to biomarkers GMD and FA. This integrated processing workflow enabled cross-modal analysis within a standardized neuroanatomical framework while preserving essential individual variation patterns.

### 2.3. Normative Model Construction and Pathological Confidence Calculation

Our DINMC framework comprises two main integrated modules: (1) Normative Model Construction, which utilizes unsupervised representation learning to construct a model that captures the normative distribution of features from large healthy cohorts, and (2) Pathological Confidence Calculation, which identifies features that are pathologically specific by computing pathological confidence values. The key procedures of the framework are outlined in Algorithm 1.

#### Algorithm 1: DINMC Framework

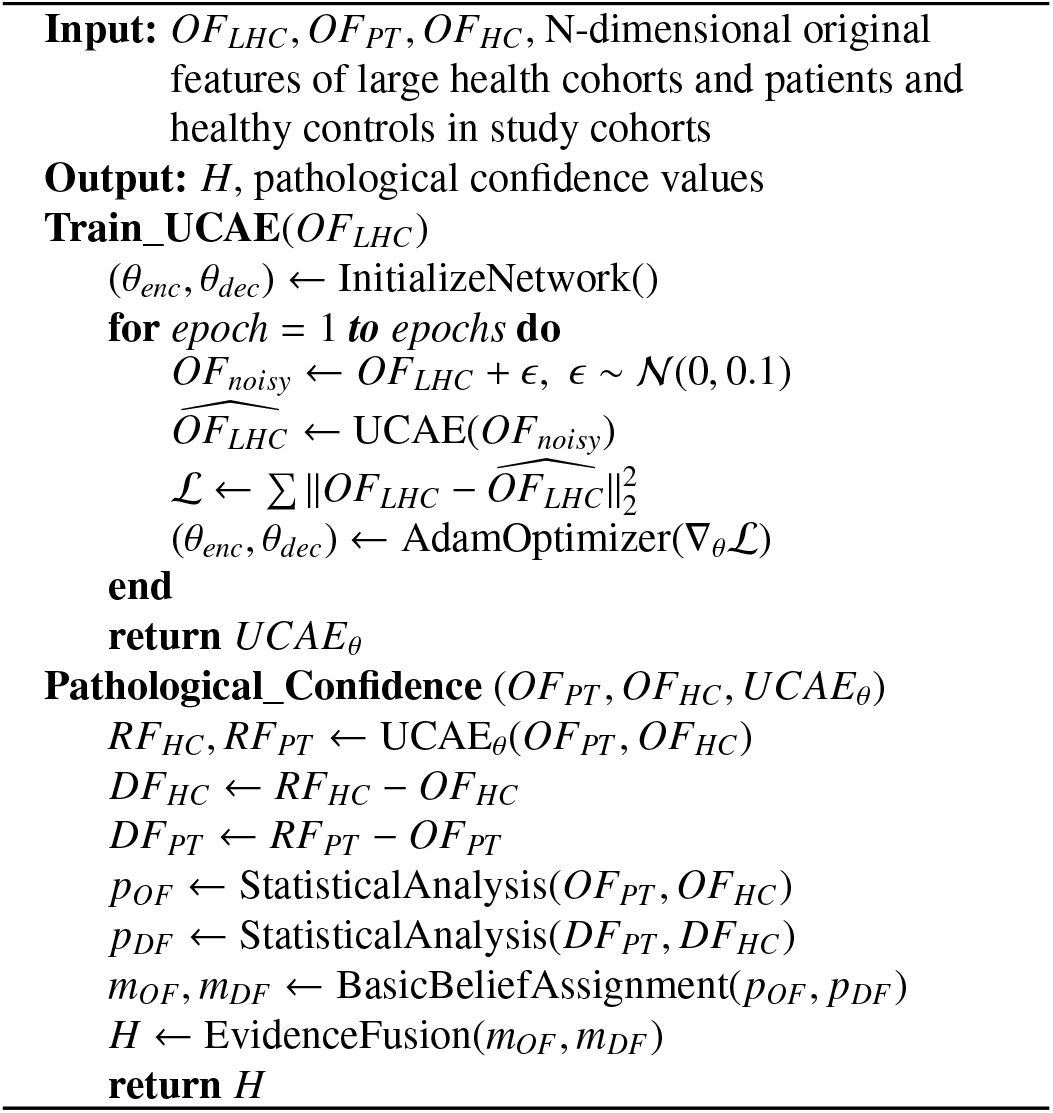

#### 2.3.1. Deep Normative Model Construction

Our normative model is a U-shaped convolutional autoen-coder (UCAE), integrating U-Net’s hierarchical feature fusion capabilities with autoencoder reconstruction principles[36]. As shown in Fig. 2, UCAE processes the multidimensional brain feature vectors through a symmetric encoder-decoder architecture optimized for precise biomarker reconstruction. This hierarchical design balances nonlinear representational capacity with geometric precision through cascaded convolutions and skip-fused upsampling, enabling context-aware recovery of normative patterns for multimodal MRI features.

**Figure 2.**
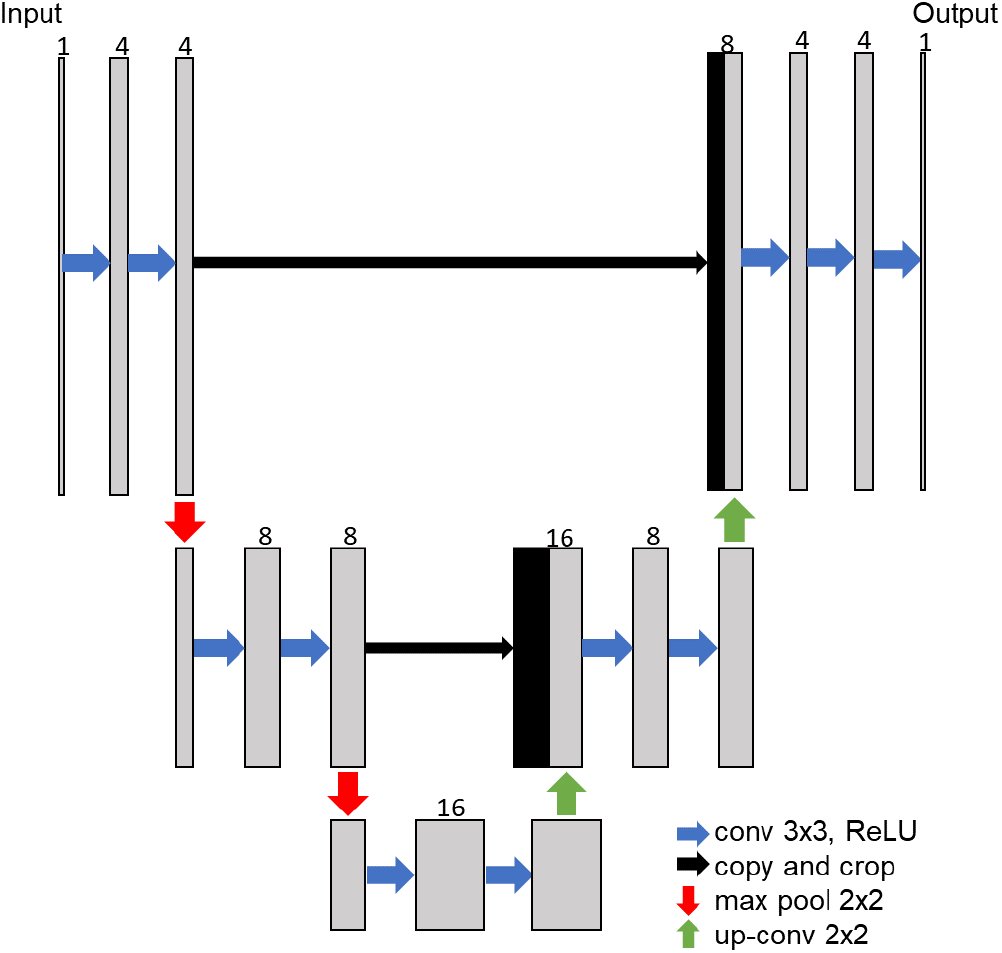
The network architecture of our U-shaped Convolutional AutoEncoder

To prevent the normative model from learning the effects of multi-site covariates during training, we applied the ComBat[18] method to harmonize the large multi-site healthy control cohorts by calibrating it to the healthy controls from the test site (CMSA). Furthermore, to learn robust normative representations, the UCAE implemented a denoising autoencoder protocol[37]. The features of large healthy cohorts were corrupted by additive Gaussian noise. The model was optimized to reconstruct the original uncorrupted features, minimizing the mean squared error (MSE) loss:

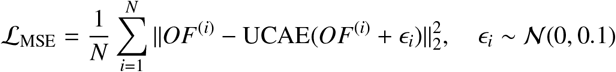

Training used the Adam optimizer[38] with scheduled learning rate decay: 0.001 (epochs 1–10), 0.0001 (epochs 11–30), 0.00001 (epochs 31–60). Optimization was performed by mini-batch gradient descent with a batch size of 20.

#### 2.3.2. Pathological Confidence Calculation

Following UCAE training in large healthy cohorts, we used this model to reconstruct the original features (OF) of both patients and healthy controls within the study cohort, resulting in reconstructed features (RF). The deviation features (DF) were derived by subtracting OF from RF. We then calculated the statistical significance for both OF and DF, obtaining p-value vectors *p*_*OF*_ and *p*_*DF*_. To integrate statistical significances from both domains, we adopted the Dempster-Shafer evidence theory framework[39], which provides a mathematical foundation for combining uncertain evidence from multiple sources. The core principles include:

1. **Frame of discernment**: Define Θ = {*A*, ¬*A*} where *A* denotes “pathology exists” and ¬*A* denotes “no pathology”
2. **Basic Belief Assignment**: Map the p-values to the belief masses *m* : 2^Θ^ → [0, 1]
3. **Evidence Fusion**: Combine *OF* and *DF* evidence by Dempster’s orthogonal rule.

##### Basic Belief Assignment

We used a negative exponential function to define a belief function *m*(*p*) to map *p*-values to the belief mass, reflecting decreasing confidence with increasing *p*-value:

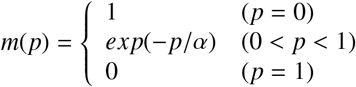

where α *alpha*is an adjustable parameter, and it is recommended to use the traditional empirical significance threshold (e.g. 0.1, 0.05, 0.01) to obtain a reasonable mapping value. This satisfies the hypothesis of the evidence theory:

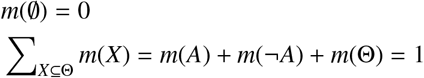

where *m*(Θ) = 1 − |*m*(*A*) − *m*(¬ *A*) |quantifies the uncertainty.

##### Evidence fusion

We fused the belief masses derived from the original (*m*_*OF*_) and deviation (*m*_*DF*_) features for each dimension to obtain the final pathological confidence *H*:

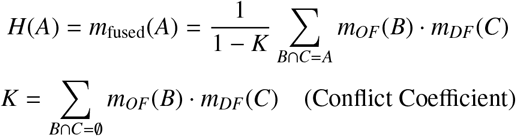

This rule combined evidence supporting pathology from both feature domains. High confidence in both domains amplified the result (e.g. *m*_*DF*_ = 0.7, *m*_*OF*_ = 0.8 ⇒ *H* ≈ 0.90), while low confidence diminished it (e.g. *m*_*DF*_ = 0.3, *m*_*OF*_ = 0.2 ⇒ *H* ≈ 0.10). Although the original and deviation features are not strictly independent, we treat them as complementary evidence sources reflecting distinct statistical perspectives (raw phenotype vs. normative deviation), following a pragmatic interpretation of evidence fusion in biomedical applications.

#### 2.3.3 Statistical Analysis

All statistical analyses were performed using custom Python code. Two-sample t-tests and Pearson correlation analyses were conducted to assess the feature differences between CMSA patients and healthy controls, as well as the correlations between features of CMSA patients and clinical scores, respectively. The parameter α controls the sensitivity of belief mass mapping and was explored across multiple values (0.1, 0.05, 0.01) to assess robustness, with α = 0.1 selected for downstream analyses based on stability considerations (see Results 3.2.1). In addition, the Benjamini–Hochberg procedure was applied to the original statistical significance *p*-values to control the false discovery rate (FDR). All statistical analyses were performed separately across three brain atlases and across GMD and FA features.

### 2.4. Prognostic Prediction

To evaluate the effectiveness of the proposed pathological confidence (*H*) in clinical prognostic prediction, we performed classification and score regression tasks after applying *H*–value based feature selection to original features. For comparison, we implemented the same models using a conventional *p*-value based feature filtering strategy as well as without any feature selection. For classification, features were selected based on statistically significant differences between patients and healthy controls, whereas for regression, features were selected based on statistically significant correlations between patient features and clinical scores. All prognostic prediction models were performed separately across three brain atlases, integrating both GMD and FA features.

#### 2.4.1. Classification

We employed a Support Vector Machine (SVM) as the classification algorithm, implemented using Python’s *sklearn.svm.SVC* with default settings [40]. To obtain low-bias estimates of classification performance on a relatively small dataset, a leave-one-out cross-validation (LOOCV) strategy was adopted. Specifically, classifiers were trained on *N* − 1 subjects and evaluated on the held-out subject, and performance metrics were subsequently averaged across all *N* folds. To prevent data leakage, all feature selection procedures were performed exclusively on the training data within each cross-validation fold. Classification performance was quantified using standardized diagnostic metrics:

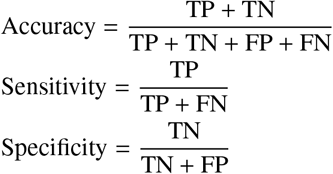

where TP = True Positives, TN = True Negatives, FP = False Positives, FN = False Negatives. The receiver operating characteristic (ROC) curves were plotted with the area under curve (AUC) values computed by trapezoidal integration. In addition, the bootstrap method was used to estimate the 95% confidence interval (CI) of AUC.

#### 2.4.2. Regression

We performed regression analyses using the Lasso regression model, implemented with Python’s *sklearn.linear_model.Lasso*. The same LOOCV and feature selection strategies as those used in the classification analysis were adopted. Model performance was quantified using the mean absolute error (MAE) and the root mean squared error (RMSE).

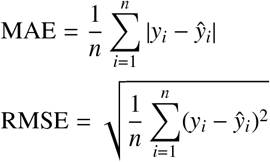

where *y*_*i*_ is the true value and *ŷ*_*i*_ is the predicted value for the *i*-th sample. In addition, the bootstrap method was employed to estimate the 95% CI for both MAE and RMSE. Furthermore, the coefficient of determination (*R*^2^) was used as an additional metric to further evaluate the predictive performance of the regression models.

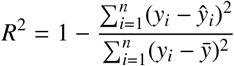

## 3. Results

### 3.1. Deep Normative Model Training

The UCAE normative model was trained for both GMD and FA features across three brain atlases (AAL116, Shen268, and AICHA384), with convergence curves displayed in Fig. 3. As shown, all models exhibited rapid convergence within the first 10 epochs, and no apparent signs of overfitting were observed throughout the training process. The inset violin plots in each panel display the MSE-based deviation metrics for CMSA patients and healthy controls. Although all models showed higher MSE in CMSA patients compared to healthy controls, most observed group-level differences did not reach statistical significance (*p* > 0.05), likely due to the limited sample size.

**Figure 3.**
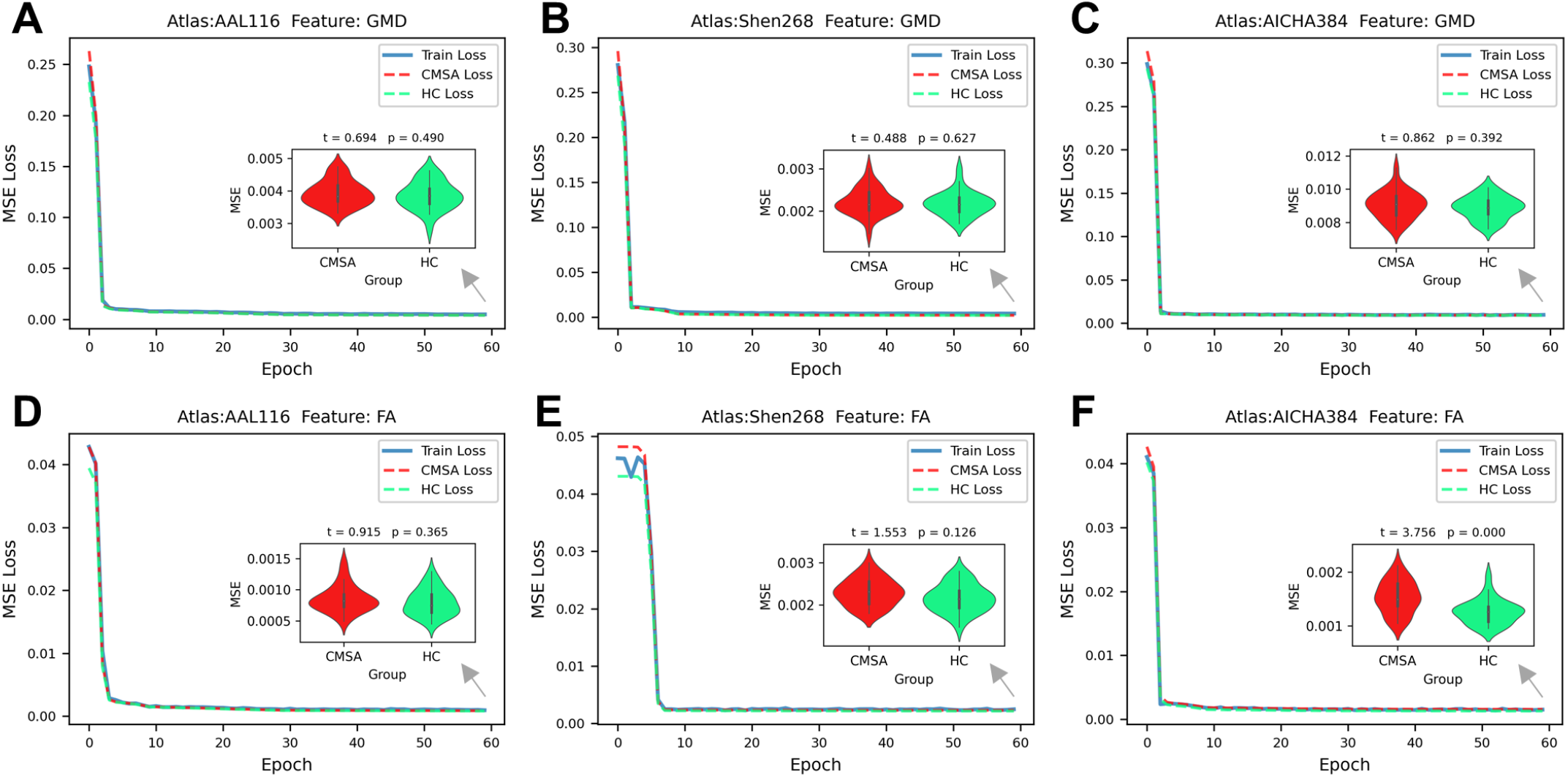
UCAE Model Training Convergence for GMD and FA Features Across Different Brain Atlases. This figure shows the convergence curves of the UCAE normative model training for GMD (A, B, C) and FA (D, E, F) features across three different brain atlases (AAL116, Shen268, and AICHA384). Each subplot represents the training loss, and test loss (CMSA loss and HC loss) over the course of 60 epochs. In each panel, a violin plot inset provides a two-sample t-test of the Mean Squared Error (MSE) between CMSA patients and healthy controls, with corresponding *t*-values and *p*-values.

These findings suggest that group-level comparisons of deviation metrics throughout the whole brain might not be sensitive enough to detect subtle differences, underscoring the importance of the brain-region-level feature-wise analyses for more accurate disease characterization.

### 3.2. Statistical Analysis

#### 3.2.1. H Distributions across α Parameters

The pathological confidence *H* is modulated by the parameter α, and we explored its distribution across three different values of α (0.1, 0.05, and 0.01). The distributions of *H* for different brain atlases and imaging features (GMD and FA) are shown in Fig. 4.

**Figure 4.**
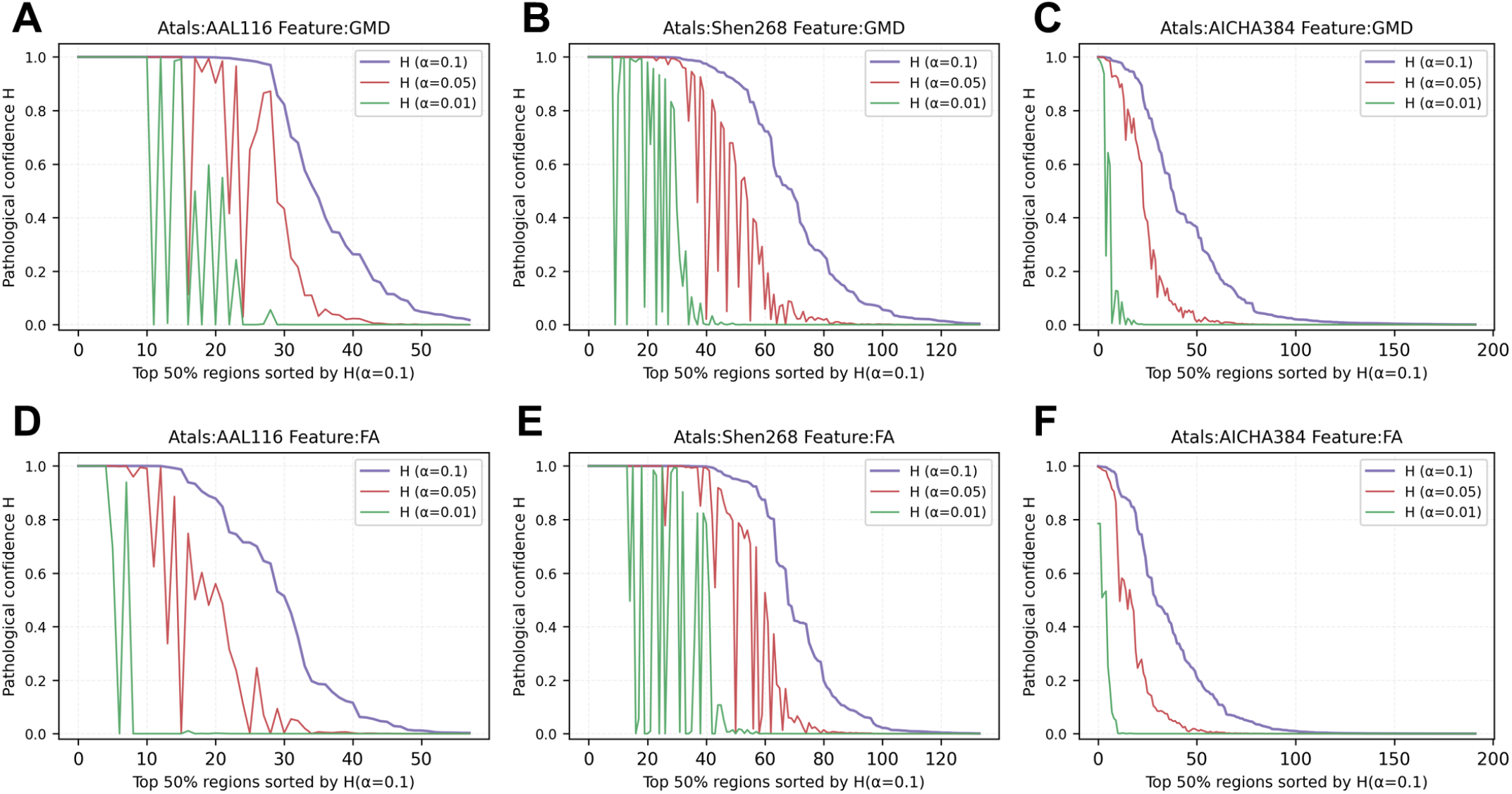
Distributions of pathological confidence *H* calculated under different parameters α across brain atlases and imaging modalities. Panels (A–C) show results for gray matter density (GMD) features using the AAL116, Shen268, and AICHA384 atlases, respectively, while panels (D–F) show corresponding results for fractional anisotropy (FA) features.

It is evident that a more lenient α leads to stronger *H* in detecting pathological changes. As α decreases, *H* gradually weakens, especially under stricter α = 0.01, where the confidence is substantially reduced. It should also be noteworthy that, under different α, the ranking of *H* for different regions is not entirely the same. This is due to the sensitivity of *H*-values to the *p*-values near the parameter α. Specifically, when *p* = α, the model assigns a relatively smaller belief mass, *m*(*p*) = *exp*(− 1) ≈ 0.37, resulting in a lower final pathological confidence value. To address this, we adopted a conventional significance threshold of *p*_*th*_ = 0.05 and set α = 0.1 for subsequent analyses, which resulted in belief masses of *m*_OF_(*p*_*th*_) = *m*_DF_(*p*_*th*_) = *exp*(−0.5) ≈ 0.6, corresponding to a pathological confidence threshold of *H*_*th*_ ≈ 0.7.

#### 3.2.2. Group-Wise Feature Analysis

The group-wise feature analysis results for GMD and FA across different brain atlases are summarized in Fig. 5, including raw *p*-values, FDR-corrected *p*-values, and *H*-values calculated with α = 0.1. After performing two-sample t-tests, no statistically significant features were observed for the AICHA384 atlas following FDR correction, whereas multiple significant features emerged for both the Shen268 and AAL116 atlases. Across the three atlases and imaging modalities, a consistent trend was observed where smaller *p*-values generally corresponded to higher *H*-values, with a few isolated exceptions. Furthermore, when considering the correspondence between pathological confidence and statistical significance, features that survived FDR correction of *p* < 0.05 showed a substantial overlap with those exceeding the pathological confidence threshold of *H* > 0.7. This highlights the substantial consistency between the proposed pathological confidence measure and conventional statistical inference methods.

**Figure 5.**
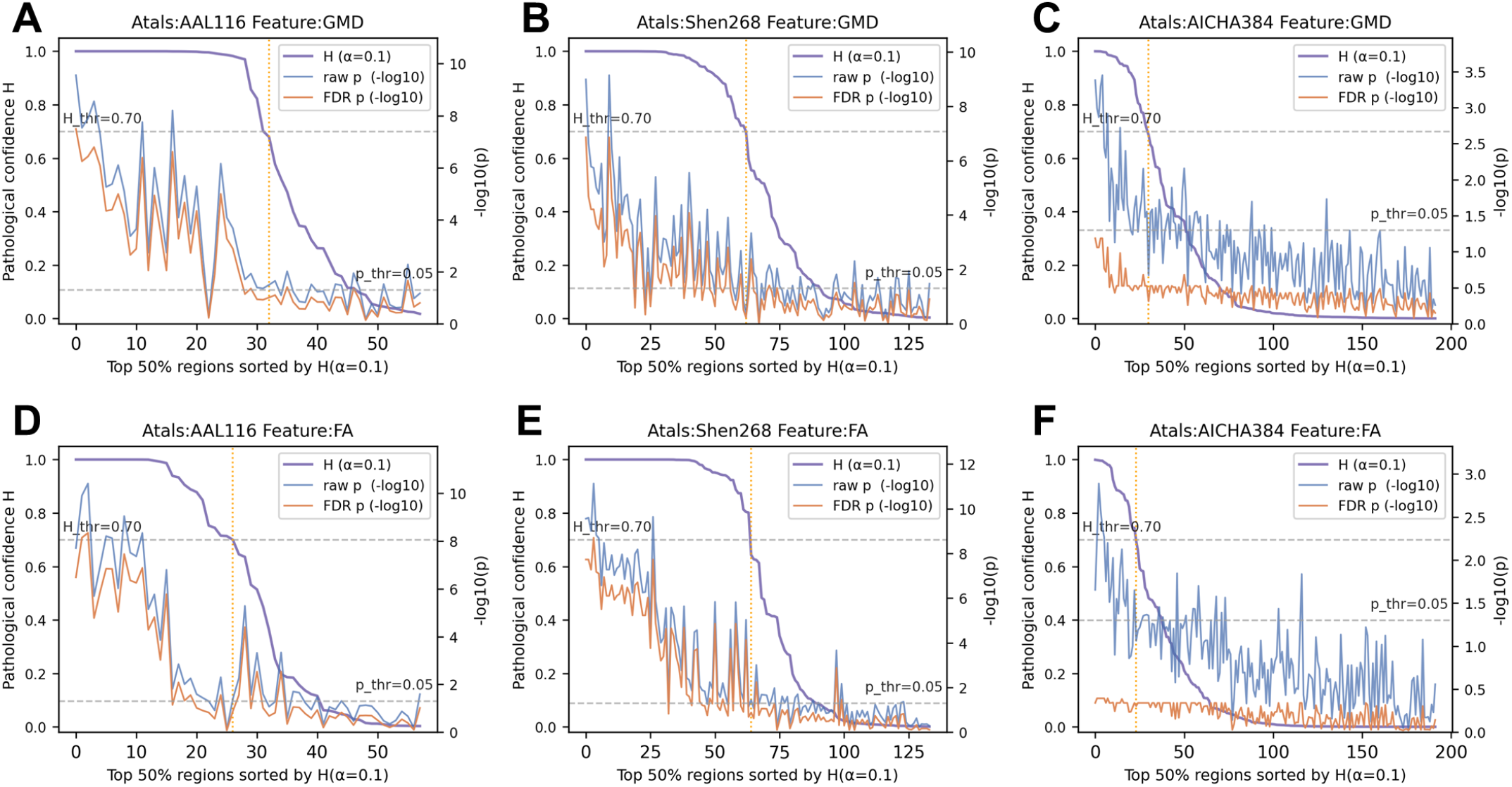
The group-wise feature analysis results for GMD and FA across different brain atlases. Panel (A,D) shows results for the AAL116 atlas, (B,E) for Shen268, (C,F) for AICHA384. Each plot compares the raw *p*-values, FDR-corrected *p*-values, and the *H*-values calculated with α = 0.1.

#### 3.2.3. Analysis of Significant Features

To further investigate the contribution of significant differences to disease pathology, we analyzed the correlation between the significantly different features and the clinical scores of the patients. Table 4 presents the number of significant features identified using FDR-corrected *p*-values and *H*-values. Specifically, the term “Sig. Diff.” denotes the features showing significant differences, while “S1 Sig.” and “S2 Sig.” correspond to these significantly different features that were significantly correlated with UMSARS-I and UMSARS-II scores, respectively. As described in 3.2.1, *p* < 0.05 and *H* > 0.7 are considered significant.

**Table 4:** Significant features identified using *p*-value and *H*-value statistics.

| Statistics | Atlas | Feature | Sig. Diff. | S1 Sig. | S2 Sig. |
| --- | --- | --- | --- | --- | --- |
| $p$-value | AAL116 | GMD | 31 | 0 | 0 |
|  |  | FA | 24 | 0 | 17 |
|  | Shen268 | GMD | 51 | 0 | 2 |
|  |  | FA | 54 | 0 | 32 |
|  | AICHA384 | GMD | 0 | 0 | 0 |
|  |  | FA | 0 | 0 | 0 |
| $H$-value | AAL116 | GMD | 32 | 0 | 3 |
|  |  | FA | 26 | 3 | 8 |
|  | Shen268 | GMD | 62 | 1 | 9 |
|  |  | FA | 64 | 4 | 12 |
|  | AICHA384 | GMD | 30 | 1 | 2 |
|  |  | FA | 23 | 0 | 1 |

From the results, it is observed that for the AAL116 and Shen268 atlases, a large number of features still show statistical significance even after FDR correction, indicating the robustness of the identified features. However, for the AICHA384 atlas, features that were initially significant disappear after FDR correction, highlighting a lack of consistent statistical support across the dataset for this particular atlas.

Interestingly, the number of significant features identified using *p*-value and *H*-value were relatively close, with both methods identifying a comparable number of features that were significant. When focusing on the clinically relevant features, both the *p*-value and *H*-value consistently demonstrated better correlations with UMSARS-II (S2 Sig.) compared to UMSARS-I (S1 Sig.). This finding suggests that the features identified as significant are more strongly related to UMSARS-II, while their relationship with UMSARS-I is comparatively weaker.

### 3.3. Prognostic Prediction

#### 3.3.1. Classification

Table 5 summarizes the classification performance for different feature selection (FS) methods across three brain atlases. Due to the small sample size, the primary evaluation metrics for classification performance were AUC and its 95% CI.

**Table 5:** Classification results for AUC, AUC CI (95%), Accuracy, Specificity, and Sensitivity across different brain atlases and feature selection (FS) methods.

| Atlas | FS Method | AUC | AUC CI (95%) | Accuracy | Specificity | Sensitivity |
| --- | --- | --- | --- | --- | --- | --- |
| AAL116 | None | 0.852 | [0.736, 0.944] | 0.821 | 0.778 | 0.862 |
|  | *p*-value Based | 0.851 | [0.726, 0.956] | 0.821 | 0.852 | 0.793 |
|  | *H*-value Based | 0.835 | [0.703, 0.946] | 0.804 | 0.852 | 0.759 |
| Shen268 | None | 0.907 | [0.797, 0.981] | 0.893 | 0.852 | 0.931 |
|  | *p*-value Based | 0.893 | [0.783, 0.981] | 0.875 | 0.815 | 0.931 |
|  | ***H*-value Based** | **0.972** | **[0.932, 0.999]** | **0.929** | **0.889** | **0.966** |
| AICHA384 | None | 0.807 | [0.671, 0.921] | 0.732 | 0.630 | 0.828 |
|  | *p*-value Based | 0.708 | [0.559, 0.837] | 0.589 | 0.593 | 0.586 |
|  | *H*-value Based | 0.738 | [0.600, 0.863] | 0.714 | 0.667 | 0.759 |

All models across the different atlases exhibited AUC values greater than 0.5, indicating that each method outperforms random classification. This confirms that there are significant feature differences between CMSA patients and healthy controls. Among the various models, the *H*-based method showed the best classification performance when using the Shen268 atlas. In this case, the *H*-based method achieved the highest AUC of 0.972, with a 95% CI ranging from 0.932 to 0.999, indicating strong and consistent predictive power.

#### 3.3.2. Regression

Table 6 presents the regression performance metrics, including RMSE, MAE, and their respective 95% CI across different methods and brain atlases. The *R*^2^ values indicate the overall effectiveness of each model in predicting the clinical scores, where a higher *R*^2^ value signifies a better predictive ability.

**Table 6:**
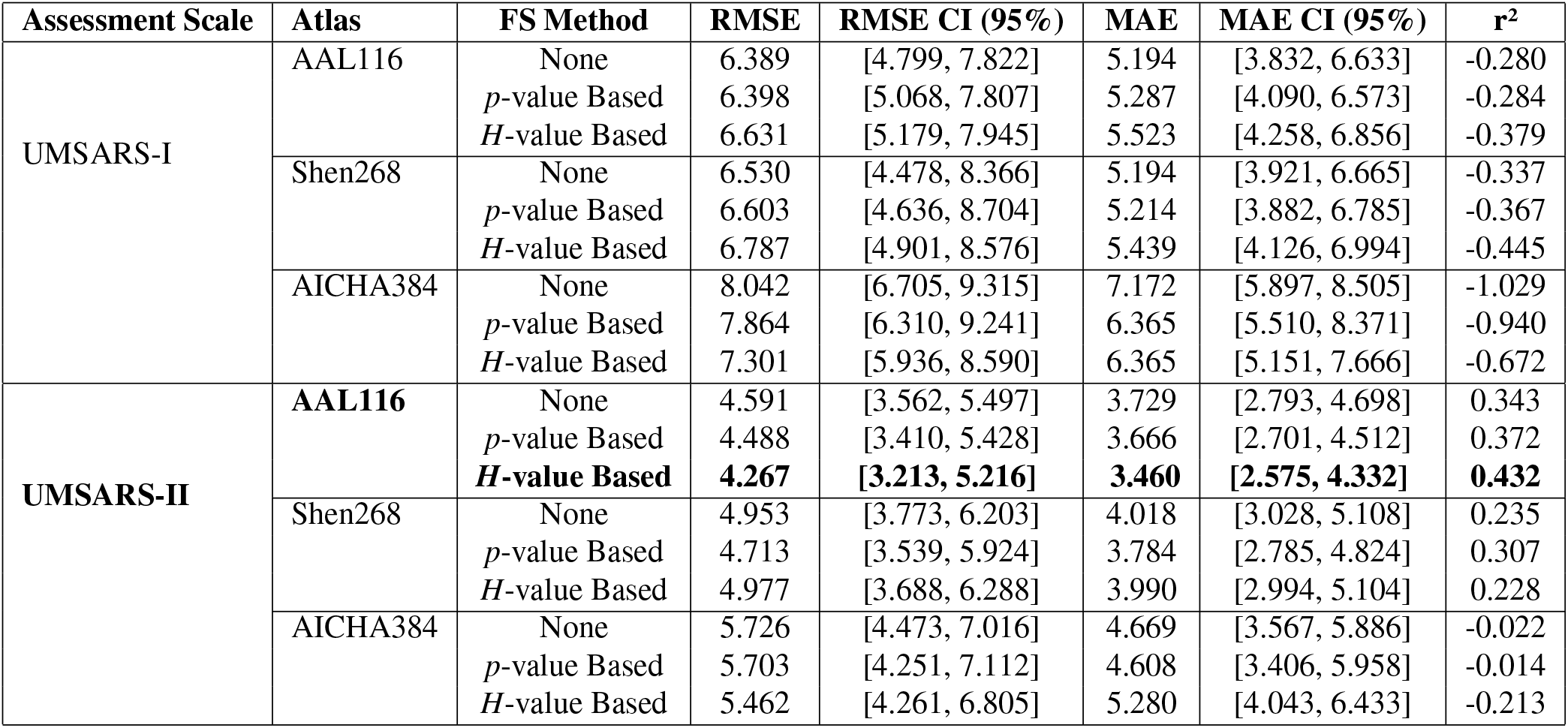
Regression results for RMSE, MAE, and *R*^2^ for UMSARS-I and UMSARS-II across different brain atlases and feature selection (FS) methods.

For all atlases, the regression results show that the *R*^2^ values for UMSARS-I are consistently negative. Negative *R*^2^ values indicate that the predictive models did not outperform a constant baseline, rather than reflecting numerical instability. This suggests that the features used in the model are insufficient or inadequately aligned with the information required to predict UMSARS-I effectively. Conversely, for UMSARS-II, the *R*^2^-values are generally positive, with the exception of the AICHA384 atlas. This indicates that the models, especially those based on UMSARS-II, perform well in predicting the clinical scores, with higher predictive power demonstrated in most atlases.

In terms of model performance, MAE and RMSE offer complementary insight into model accuracy and error. Among the various models, the *H*-based method showed the best regression performance when using the AAL116 atlas and preditcing UMSARS-II. Specifically, this model achieved the lowest RMSE of 4.267 with a 95% CI ranging from 3.213 to 5.216 and the lowest MAE of 3.460 with a 95% CI ranging from 2.575 to 4.332, indicating strong predictive power.

#### 3.3.3. Summary

Overall, these results underscore the influence of different brain atlases on model performance. The AAL116 and Shen268 atlases respectively provided the best results, particularly with the *H*-based method, while the performance of the AICHA384 atlas was consistently the worst. This highlights the *H*-based method’s ability to capture the pathological alterations of CMSA and emphasizes the importance of selecting the appropriate brain atlas for optimal prediction results.

## 4. Discussion

In this study, we proposed the DINMC framework, which consists of two integrated components. The first, *Deep Normative Model Construction*, uses multi-site healthy data to train the UCAE normative model, which then reconstructs the original features of both patients and controls within the study cohort, generating deviation features. The second component, *Pathological Confidence Calculation*, quantifies the pathological relevance of these features by calculating pathological confidence values using Dempster-Shafer evidence theory. Our results demonstrate that the pathological confidence patterns computed using the DINMC framework provide a more accurate reflection of the pathological progression of CMSA compared to traditional statistical approaches based on significance thresholds. Furthermore, feature selection based on pathological confidence thresholds outperformed traditional *p*-value threshold-based methods in clinical prognosis prediction tasks, including classification and regression.

### 4.1. Interpretability of Normative Models in Neuroimaging

Neuroimaging features are highly susceptible to multi-site effects caused by differences in scanner manufacturers and acquisition protocols, which can bias normative model estimations unless accounted for [41, 16]. While some harmonization techniques help, they may also remove meaningful biological signals [42, 43]. Although traditional case-control studies that use the same equipment for both patients and healthy controls can mitigate some of these effects, such studies are often limited by small sample sizes. Existing normative studies attempt to address these challenges by using larger samples and innovative methodologies, but no approach has yet fully resolved the issues related to multi-site covariates. In particular, when applying a model to data from a previously unseen site, a certain amount of data from that site is still required for fine-tuning or calibration.

Our DINMC framework does not seek to resolve multi-site issues from a theoretical perspective but rather focuses on improving the interpretability of normative models in neuroimaging applications. Existing normative models often rely on feature deviations as standalone substitutes for original features, but lack statistical assurances that these deviations truly reflect pathology. Moreover, normative models built with large samples often focus on relatively simple structural features, such as brain development, limiting their clinical applicability. In contrast, our approach integrates the interpretability of hypothesis testing frameworks, combining statistical results from both the original and deviation feature spaces. By framing statistical differences in terms of pathological confidence, our model shifts the focus from controlling error rates to understanding the biological plausibility of feature deviations, providing more robust interpretations of the results. This bridges the gap in existing normative models, ensuring that deviation features are not simply a replacement for original features, but offer meaningful insights into disease pathology.

### 4.2. Impact of Brain Parcellation and Atlas Selection

The impact of preprocessing on neuroimaging research is significant, and brain parcellation is a critical preprocessing step in neuroimaging studies[44]. Using brain atlases to define a unified space and “node” structure offers at least two advantages.

First, Using brain atlases creates standardized feature spaces that support cross-modal and cross-study comparability [45, 46]. Currently, neuroimaging studies use varying coordinate systems, resolutions, and statistical methods, which can hinder the reproducibility and integration of results. By using a common feature space, brain atlases facilitate the fusion of multimodal data and are particularly important for building functional normative models in future studies.

Second, parcellation also reduces the burden of multiple comparisons and enhances signal-to-noise ratio compared to voxel-based analyses [47]. When voxel-based features are analyzed, the number of statistical tests can reach millions, and traditional methods such as FDR correction often become overly conservative, potentially masking meaningful biological effects. This has led to the development of specialized methods, such as network-based statistics [48] or cluster-based statistics [49], but these also limit the reproducibility and integration of results. In contrast, averaging features across the entire brain results in a low signal-to-noise ratio and weak interpretability. For instance, when analyzing the MSE deviation metrics for CMSA patients and healthy controls across the whole brain, although there was a trend of higher error in patients, the difference was not statistically significant and could not be directly applied.

In this study, we used three different brain atlases: AAL116, Shen268, and AICHA384. The AAL116 atlas is based on anatomical structures, including both the brain and cerebellum, while the Shen268 and AICHA384 atlases use functional connectivity to define brain regions. The AICHA384 atlas only includes brain regions. Our statistical analysis revealed that the significance of original p-values for features based on the AICHA384 atlas was weaker than that computed for the same features in the other two atlases. As a result, after FDR correction, no features in the AICHA384 atlas reached statistical significance.

### 4.3. Analysis of CMSA Pathology

CMSA is a progressive neurodegenerative disorder characterized by α-synuclein-positive glial inclusions that primarily affect cerebellar and olivopontine circuits [50]. Previous studies [30] have confirmed widespread GMD atrophy and FA reduction in cerebellar subregions using the AAL116 atlas. In our study, we observed that the AICHA384 atlas, which does not include cerebellar regions, exhibited the worst performance in both statistical analysis and prognostic prediction. This observation is highly consistent with the known cerebellar-dominant pathology of CMSA, reinforcing the biological validity of atlas-dependent findings. This finding highlights the crucial role of the cerebellum in the pathological progression of CMSA. The absence of cerebellar structures in the AICHA384 atlas likely limits the model’s ability to capture the disease-related abnormalities in these regions, which are key to understanding the clinical manifestations of CMSA.

Furthermore, we observed that the model’s performance for predicting UMSARS-II scores was consistently better than for UMSARS-I. The higher accuracy of UMSARS-II prediction may be attributed to the fact that UMSARS-II reflects more direct motor dysfunction, which is often more pronounced and easier to correlate with neuroimaging features. In contrast, UMSARS-I focuses on activities of daily living, which can be influenced by factors such as cognitive or mood disturbances, leading to more variability in the assessment and potentially making it more challenging to predict accurately using neuroimaging data. Therefore, the stronger correlation between neuroimaging features and UMSARS-II scores further reinforces the notion that motor impairments in CMSA are more directly linked to structural brain changes than functional limitations assessed in UMSARS-I.

### 4.4. Limitations and Future Directions

Several limitations should be considered in this study. First, the theoretical foundation of our pathological confidence calculation requires further refinement. The belief function currently relies on empirically determined parameters, which could introduce bias and impact the robustness of our findings. Second, the Dempster-Shafer framework assumes that the evidence sources are statistically independent, a condition not fully satisfied in our methodology. Additionally, the current validation is limited to structural neuroimaging features, and future studies should include functional metrics, which are crucial to understanding brain organization and offer a more holistic perspective. Finally, longitudinal studies are needed to capture the dynamic pathological progression of CMSA, enabling a better understanding of temporal changes in biomarkers.

## 6. Conclusion

This study introduces an interpretable normative modeling framework that integrates statistical hypothesis testing with deep autoencoder-based reconstruction. Validated on a cerebellar-type multiple system atrophy cohort, the proposed DINMC framework enables the quantification of pathological confidence at the feature dimension level, providing biologically grounded interpretation of neuroimaging deviations. The results demonstrate that confidence-guided feature selection improves downstream classification and clinically relevant score prediction, particularly for motor impairment. The framework is generalizable to other multimodal neuroimaging settings and offers a scalable paradigm for interpretable precision analysis in biomedical engineering.

## Ethics statement

The study was approved by the Medical Research Ethical Committee of AeroSpace Center Hospital (No.2023-071) in accordance with the Declaration of Helsinki and relevant ethical guidelines. Written informed consent was obtained from each participant or their legal representatives.

## CRediT authorship contribution statement

**Zewu Ge:** Conceptualization, Data curation, Formal analysis, Investigation, Methodology, Software, Visualization, Writing – original draft, Writing – review & editing. **Shui Liu:** Investigation, Resources, Data curation, Writing – review & editing.**Weibei Dou:** Conceptualization, Resources, Funding acquisition, Project administration, Resources, Supervision, Validation, Writing – review & editing.

## Acknowledgements

This work was supported by the National Key Research and Development Program of China under Grant 2022YFC3601100 and Grant 2022YFC3601105. Zewu Ge wishes to thank Dong-fang Hospital of Beijing University of Traditional Chinese Medicine for the data collection of the CMSA cohort. Research reported in this publication was supported by the National Institute On Aging of the National Institutes of Health under Award Number U01AG052564. The content is solely the responsibility of the authors and does not necessarily represent the official views of the National Institutes of Health.

